# Computational drug repositioning of bortezomib to reverse metastatic effect of *GALNT14* in lung cancer

**DOI:** 10.1101/394163

**Authors:** Ok-Seon Kwon, Haeseung Lee, Hyeon-Joon Kong, Ji Eun Park, Wooin Lee, Seungmin Kang, Mirang Kim, Wankyu Kim, Hyuk-Jin Cha

## Abstract

Although many molecular targets for cancer therapy have been discovered, they often show poor druggability, which is a major obstacle to develop targeted drugs. As an alternative route to drug discovery, we adopted an *in silico* drug repositioning (*in silico* DR) approach based on large-scale gene expression signatures, with the goal of identifying inhibitors of lung cancer metastasis. Our analysis of clinicogenomic data identified GALNT14, an enzyme involved in O-linked N-acetyl galactosamine glycosylation, as a putative driver of lung cancer metastasis leading to poor survival. To overcome the poor druggability of GALNT14, we leveraged Connectivity Map approach, an *in silico* screening for drugs that are likely to revert the metastatic expression patterns. It leads to identification of bortezomib (BTZ) as a potent metastatic inhibitor, bypassing direct inhibition of poorly druggable target, GALNT14. The anti-metastatic effect of BTZ was verified *in vitro* and *in vivo*. Notably, both BTZ treatment and *GALNT14* knockdown attenuated TGFβ-mediated gene expression and suppressed TGFβ-dependent metastatic genes, suggesting that BTZ acts by modulating TGFβ signalingTaken together, these results demonstrate that our *in silico* DR approach is a viable strategy to identify a candidate drug for undruggable targets, and to uncover its underlying mechanisms.

## Introduction

In the context of personalized anti-cancer therapy based on targeting specific proteins with the goal of lowering cancer-related mortality (1), a great deal of effort has been devoted to identifying both molecular targets and accompanying drugs (2, 3). However, the fraction of patients eligible for personalized anti-cancer therapy is very limited (4) due to the poor druggability of newly identified molecular targets, notwithstanding recent advances in strategies in drugging ‘undruggable’ proteins (5, 6).

An alternative approach to matching candidate drugs to poorly druggable cancer targets is *in silico* drug repositioning (*in silico* DR) (7, 8). Due to the well-characterized pharmacology and safety of approved drug libraries(9), this approach has the potential to reduce cost and attrition during the clinical phases of drug development. Several approaches to DR have been tested in the context of oncology (10) and a few of the resultant drugs, including celecoxib and thalidomide, have been approved by the FDA for repurposing as anti-cancer therapeutics (11). Along with recent advances in sequencing technologies, chemogenomic databases containing drug-induced gene expression profiles provide clues regarding potential treatments for personalized cancer targets, and can also suggest candidate drugs based on tailored gene signatures of cancers upon identification of molecular targets (12). The Connectivity Map (CMap), a collection of genome-wide expression profiles of cell lines treated with >20,000 chemicals(13), has been used to identify candidate drugs for certain cancer types (14–16).

N-acetyl-galactosaminyltransferases (GalNAc-Ts or GALNTs) are key enzymes that initiate O-linked N-acetyl galactosamine (GalNAc) glycosylation. This process is an important step in the synthesis of Thomsen-nouvelle (Tn) antigens, which are well-characterized tumor-associated molecules (17). In particular, *GALNT14* has been examined in the context of apoptotic signaling (18, 19); invasion and migration of breast (20, 21), ovary (22), and lung (23) cancers; and multi-drug resistance of breast cancer cells (24). Moreover, *GALNT14* expression is not only a prognostic marker in neuroblastoma (25) and lung cancer (23), but also a predictive marker for Apo2L/TRAIL-based cancer therapy (18), although a randomized phase II study based on the predictive marker of GALNT14 for dulanermin did not improve patient outcome (26).

In this study, through transcriptome analysis of the TCGA dataset and *in vitro* and *in vivo* studies, we demonstrated that *GALNT14* is strongly associated with lung cancer recurrence due to the high migration and invasion properties of tumor cells. Rather than attempting direct inhibition of the poorly druggable GALNT14 protein or downstream signaling, we leveraged large-scale drug-induced transcriptome data to identify candidate drug(s) likely to reverse *GALNT14*-dependent gene expression, *i.e*., drugs that led to transcriptomic changes similar to those induced by *GALNT14* depletion. We successfully identified an anti-metastatic candidate drug that mimicked *GALNT14* depletion. The results demonstrate that this approach represents a viable strategy for discovering candidate drugs for many other undruggable targets.

## Results and discussion

### *GALNT14* as a putative molecular target for lung cancer metastasis

Identification of molecular targets in recurrent cancers is essential not only for predicting prognosis, but also for matching specific drug–target pairs if they are available. To identify potential molecular targets related to cancer recurrence, we assembled transcriptome data and clinical information from 516 lung cancer patients from the TCGA LUAD cohort (Figure. 1A). Concentrating on molecular targets relevant to recurrent lung cancer, we performed a series of relapse-free survival (RFS) analyses and differential expression analyses. Expression of seven genes (*GALNT14, COL7A1, GPR115, C1QTNF6, KRT16, INHA*, and *TNFSF11*) was significantly associated with cancer progression, recurrence (Figure. 1B), and overall survival (Figure. S1A), indicating that these genes are potentially valuable as predictors of poor prognosis. Notably, metastasis-related genes were significantly overrepresented in gene lists selected by both RFS (*P* = 2.3 × 10^−6^) and differential expression (*P* = 5.2 × 10^−19^) analysis, including two genes (*e.g., TNFSF11* and *INHA*) among the seven aforementioned candidates. To further confirm the relevance of each gene to metastasis or tumorigenesis, we divided lung cancer patients into two groups (low or high) using the median expression of each gene as the cutoff. Both metastasis and tumor signatures were positively enriched in the high-expression groups of all seven genes (Figure. 1C), and the significant enrichment was observed for *GALNT14* (Figure. 1D).

**Figure 1.**
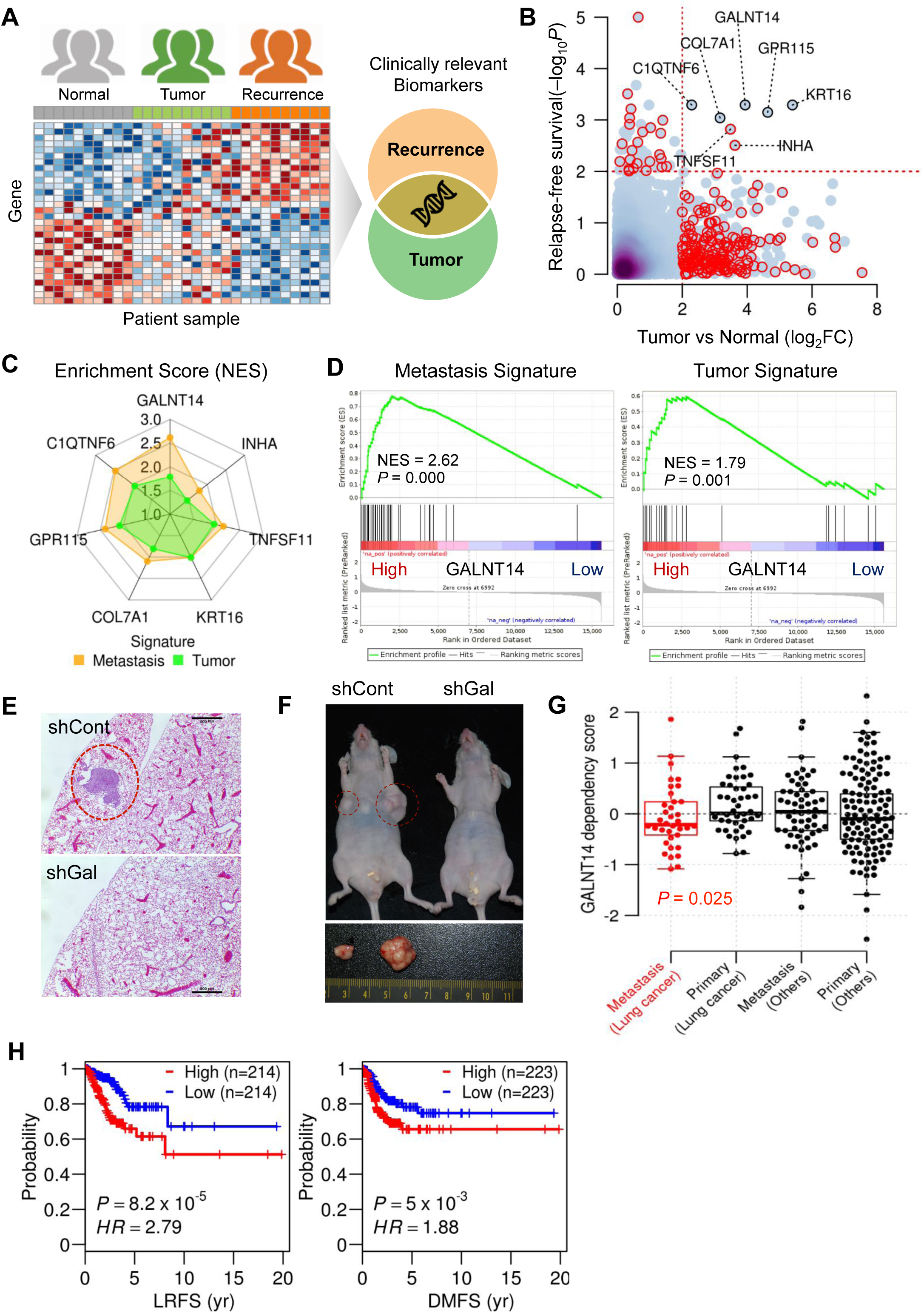
*GALNT14* as a putative molecular target for lung cancer metastasis. **A.** Identification of therapeutic targets for both cancer progression and recurrence based on transcriptomic analysis of lung cancer patients. **B.** Selection of therapeutic target candidates that are differential expressed between tumor and normal tissue as well as significantly associated to RFS in the 516 patient dataset from TCGA LUAD. Red circles indicate metastasis-related genes annotated by MSigDB. **C.** Enrichment analysis of metastatic and tumor signatures between high- and low-expressed patient groups for each of the seven candidate genes. **D.** The normalized enrichment score (NES) was calculated by Gene Set Enrichment Analysis (GSEA) for the metastasis (left) and tumorigenesis signature (right) after all the genes were ranked by their expression fold change. **E-F.** Control H460 (shCont) and GALNT14 knockdown (shGal) cells were injected into lateral tail vein (E) or flanks (F) of nude mice. The representative H&E staining images of tumor-bearing lung (E) and tumors (F) were presented. **G.** Comparison of *GALNT14* dependencies among metastatic and primary cell lines from lung and other types of cancer from the *Project Achilles* dataset. Only metastatic lung cells show a significant dependency on *GALNT14*. **H.** Comparison between the high- and low-expression group for *GALNT14* in terms of locoregional recurrence-free survival (LRFS), and distant metastasis-free survival (DMFS) in LUAD patients.

*GALNT14*, which encodes a glycosyltransferase involved in O-glycosylation, has been implicated in both tumor malignancy (25) and metastasis (20, 21, 23). As expected, metastatic (Figure. 1E) and tumorigenic potentials (Figure. 1F) were markedly attenuated by loss of *GALNT14*, indicating that the gene is important for metastasis as well as tumorigenesis. Further, metastatic lung cancer cells were more vulnerable to *GALNT14* depletion than non-metastatic or other types of cancers in the *Project Achilles* dataset(27), a genome-scale RNAi screening data from for 501 cancer cell lines, including 126 cell lines originating from metastatic patients (Figure. 1G). Other candidate genes were less vulnerable in metastatic lung cancer (Figure. S1B). Consistent with this, *GALNT14* expression shows a clearly negative correlation with both locoregional recurrence-free survival (LRFS) and distant metastasis-free survival (DMFS) (Figure. 1H) as well as overall survival (Figure. S1A) in the TCGA LUAD cohorts. In addition, normal lung expresses only a low level of *GALNT14*, and there is a large gap between normal and lung cancer tissue (Figure. S1C)(28). All these results suggested *GALNT14* as a promising molecular target for lung cancer metastasis to improve patient survival.

### Computational repositioning of BTZ to reverse the *GALNT14* expression signature

Although *GALNT14* may be a potential therapeutic target for metastatic lung cancer, GALNTs remain poorly druggable despite several attempts to find specific inhibitors (29, 30). Notably in this regard, GALNT14-dependent metastatic potential is governed by induction of transcription factors (e.g. *HOXB9* or *SOX4*) rather than by altered glycosylation (20, 23). Therefore, rather than inhibiting GALNT14 directly, we leveraged CMap dataset to virtually screen drugs that mimic the effects of *GALNT14* depletion at the transcriptome level. In particular, we focused on genes associated with metastasis or *GALNT14* expression, and considered that these genes should be suppressed in order to restore the metastatic to the normal. To this end, we defined two distinct *GALNT14* signatures. We collected a comprehensive list of 3711 metastasis-related genes and selected two sets of genes among them: (i) 20 genes up-regulated in the *GALNT14*-high group in the TCGA LUAD cohort, and (ii) 49 genes down-regulated by *GALNT14* knockdown in the H460 cell line. Accordingly, our subsequent predictions using *GALNT14* signatures would prioritize drugs that are probably relevant to both GALNT14-dependence and metastasis.

Using the two *GALNT14* signatures, we performed two independent predictions by CMap analyses (Figure. 2A). Candidate drugs were prioritized according to their DR scores (See Materials and Methods in detail). Two drugs, dexamethasone (DEX, an anti-inflammatory corticosteroid) and bortezomib (BTZ, a first-in-class proteasome inhibitor used to treat multiple myeloma), were among the top candidates in both predictions (Figure. 2B). We then validated expression levels of *SOX4, AREG*, and *VCAN*, which are strongly associated with metastasis and regulated by *GALNT14* (20, 23) (Figures. S2B and S2C). Among genes differentially expressed in response to either DEX or BTZ in the CMap dataset, *SOX4* (but not *AREG* or *VCAN*) was commonly altered, indicating that *SOX4* could serve as a validation marker (Figure. S2C). As predicted, we observed dose-dependent suppression of *SOX4* (but not *VCAN*) in H460 cells treated with either DEX or BTZ. Although BTZ suppressed *SOX4* less effectively than DEX (EC_50_: 15 nM *vs*. 5 nM, respectively) (Figure. S2D), BTZ treatment led to a significant reduction in the migration capacity of H460 cells, whereas DEX did not (Figure. 2C). The EC50 for proteasome inhibition was around 20 nM (Figure. 2D and S2E), a concentration at which BTZ clearly inhibited migration (Figures. 2E and S2F) and invasion (Figure. 2F) by lung cancer cells, while it did not affect *GALNT14* expression (Figure. S2G), cell viability (Figures. 2G and S2H) nor the cell proliferation (Figures. 2H and S2I). Of note, A549 with relatively lower GALNT14 expression than H1299 and H460 (Figure. S2J) did not respond to BTZ treatment in migration (Figure. S2K) and invasion (Figure. S2L), while proteasome inhibition by BTZ occurred (Figure. S2M), suggesting that anti-invasion/migration effect of BTZ would be associated with GALNT14 expression.

**Figure 2.**
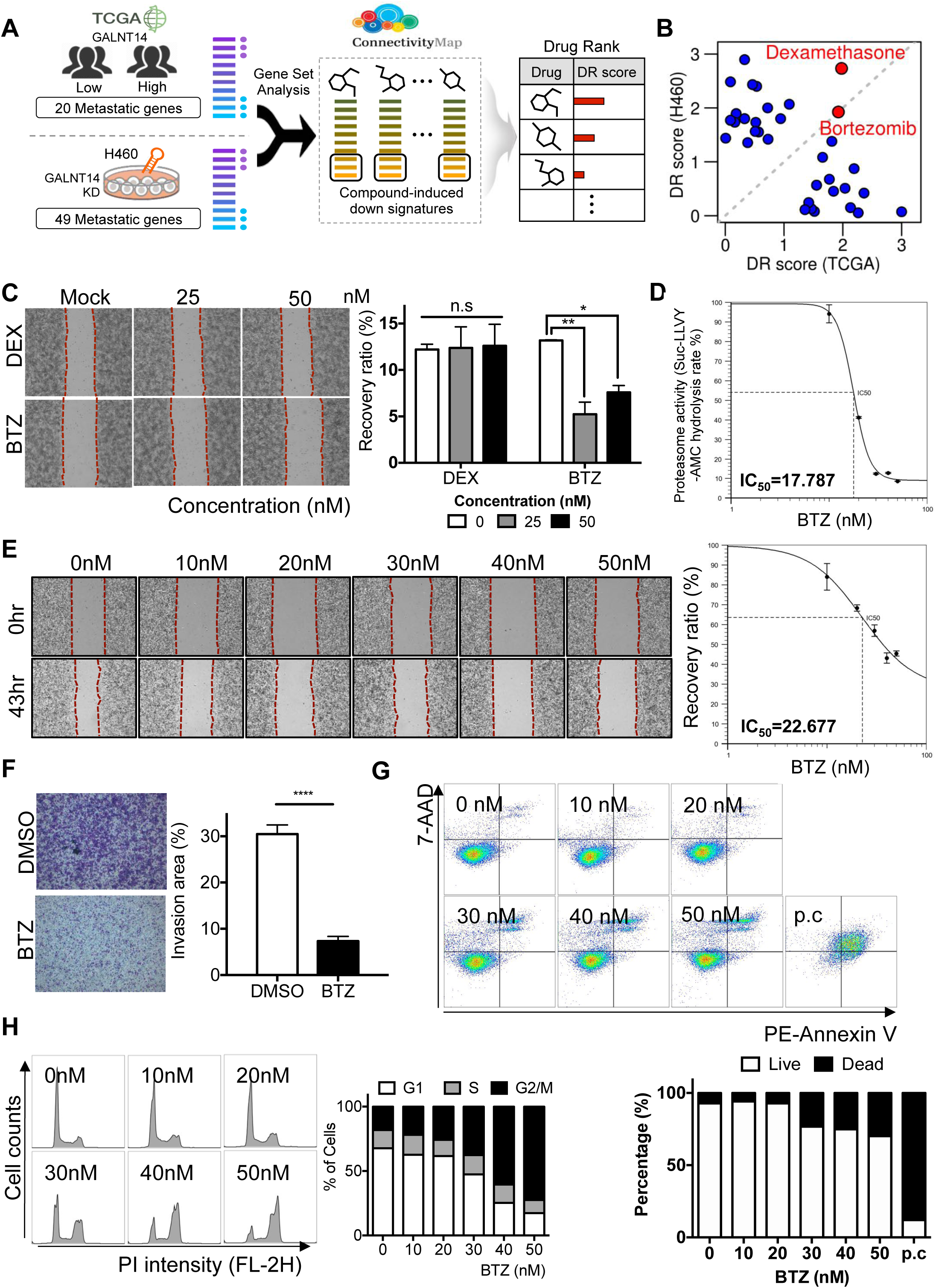
Computational repositioning of BTZ to reverse the *GALNT14* expression signature. **A.** CMap analyses to prioritize anti-metastatic candidate drugs using the two independent *GALNT14* signatures. Candidate drugs were prioritized according to the similarities between drug-induced down-DEGs and the two *GALNT14* signatures. **B.** DR score of top candidate drugs selected by the *GALNT14* signatures from TCGA (x-axis) and H460 (y-axis). The two drugs marked in red (dexamethasone and bortezomib) got high scores in both predictions. **C.** Representative microscopic images of cell migration after treatment of indicated dose of (nM) of BTZ and DEX (left), Relative recovery ratio of indicated group was graphically presented. (right, n.s: not significant) **D.** Proteasome activity in H460 cells after indicated dose of BTZ was graphically presented. **E.** Representative microscopic images of cell migration at 43 hours after treatment of indicated dose of (nM) of BTZ (left), Relative recovery ratio of indicated dose was graphically presented (right) **F.** Representative image of invaded cell through two-chamber model after DMSO or 20 nM of BTZ treatment (left), Relative invasion area was graphically presented (right). **G.** Flow cytometry for Annexin V and 7-AAD at 24 hours after indicated dose of BTZ (top), Graphical presentation of live (white box) and dead (black box) was shown (bottom). **H.** Flow cytometry of cell cycle profile at 24 hours after indicated dose of BTZ treatment (left), Graphical presentation of each cell cycle (G1, S and G2/M) were shown (right).

### The effect of BTZ is independent of proteasome inhibition

Given that the anti-migration/invasion effect of BTZ occurred at a concentration that also inhibited proteasome activity (Figures. 2D and E), we sought to determine whether this effect was a result of proteasome inhibition *per se*. To investigate this issue, we first compared the chemical structure of three FDA-approved proteasome inhibitors, BTZ, carfilzomib (CFZ), and ixazomib (IXZ), all of which are approved for treatment of multiple myeloma (31). CFZ displayed similar profiles with regards to Tanimoto coefficients or Jaccard index of molecular fingerprints (32) (Figure. 3A). In contrast to BTZ, CFZ also inhibited proteasome activity (Figure. S3A) and stabilized well-characterized proteasome targets β-catenin, Cyclin D1, and p27 (Figure. 3B), but failed to suppress migration (Figure. 3C) and invasion (Figure. 3D) at the same concentration. Moreover, the boronic acid moiety responsible for the proteasome inhibition (33) was present in both BTZ and IXZ. However, like CFZ, treatment with IXZ could not inhibit migration (Figure. S3B). These data suggest that BTZ has an off-target effect that is independent of proteasome inhibition.

**Figure 3.**
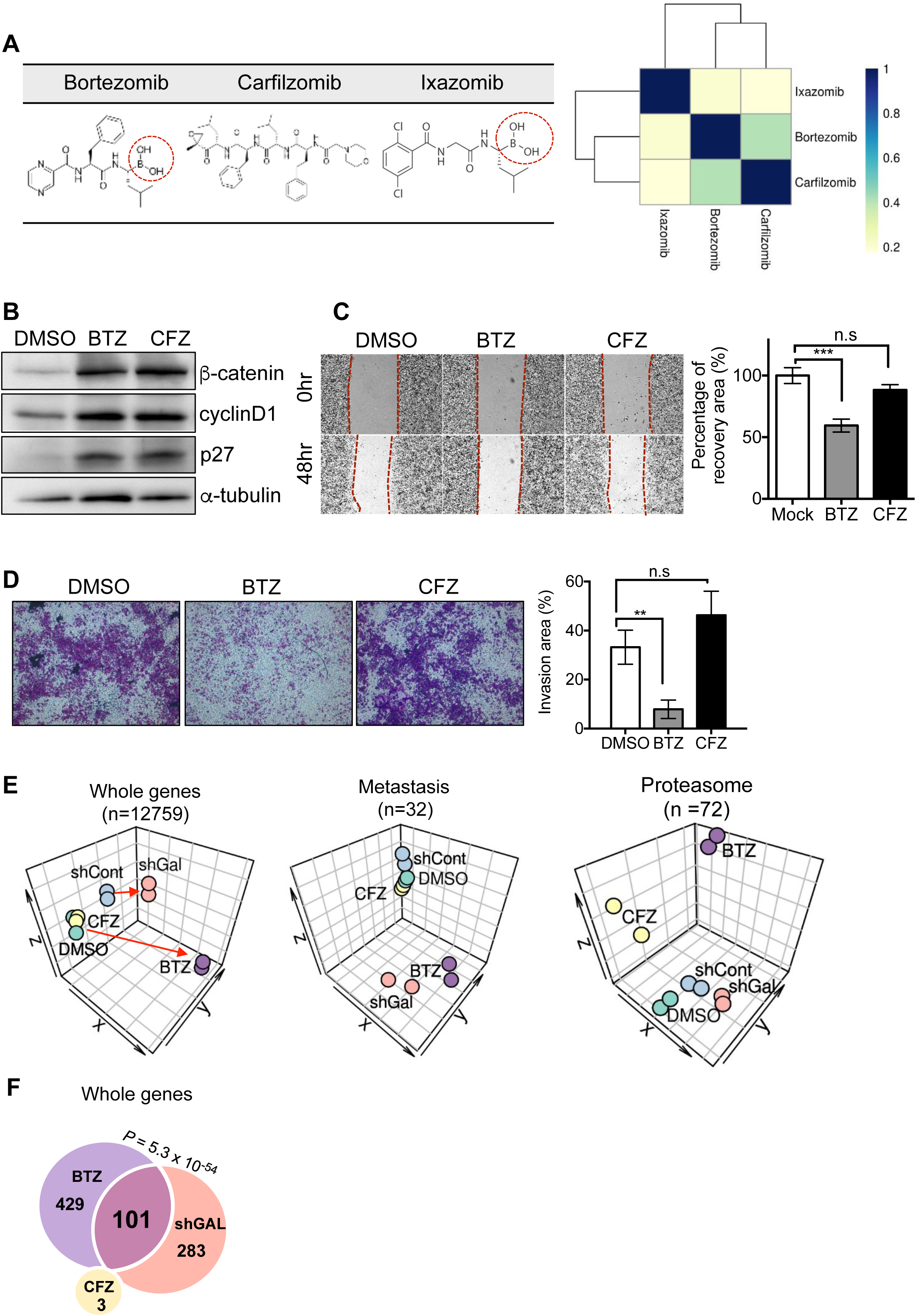
The effect of BTZ is independent of proteasome inhibition. **A.** Chemical structure of proteasome inhibitors (bortezomib, carfilzomib, and ixazomib) and their Tanimoto similarity heatmap. Red circle indicates the boronic acid structure. **B.** Immunoblotting for β-catenin, Cyclin D1 and p27 after treatment of BTZ (20 nM) and CFZ (20 nM), α-tubulin for equal protein loading control **C.** Representative microscopic images of cell migration at 43 hours after 20 nM of BTZ and CFZ treatment (left), Relative recovery ratio was graphically presented (right, n.s: not significant). **D.** Representative image of invaded cell through two-chamber model after DMSO or 20 nM of BTZ or CFZ treatment (left), Relative invasion area was graphically presented (right, n.s: not significant). **E.** Sample clustering using t-SNE based on the expression of whole genes, metastasis-related genes and proteasome-related genes. Each sample was colored according to its perturbation type. **F.** Venn diagram of differentially down-regulated genes by BTZ, CFZ and shGAL perturbation in the gene space of whole genes.

To confirm that the anti-migration/invasion effect of BTZ is not dependent on proteasome inhibition, we compared transcriptome profiles of H460 cells treated with BTZ, CFZ and depleted of *GALNT14* (sh*GAL*). Perturbation by BTZ and sh*GAL* (but not CFZ) induced similar transcriptomic changes relative to the control (Figure. 3E, left panel). Moreover, the expression patterns of metastatic signature genes were even more similar between BTZ and sh*GAL*, whereas CFZ had a minimal effect on only a few genes (Figure. 3E, middle panel). By contrast, expression of proteasome-related genes was altered significantly by both drugs, but only marginally by sh*GAL* (Figure. 3E, right panel). Genes down-regulated by BTZ and sh*GAL* overlapped significantly (101 common genes; *P* = 5.3 × 10^−54^), suggesting that BTZ treatment partially mimics depletion of *GALNT14* (Figure. 3F). These results are consistent with the phenotypic outcomes, i.e., BTZ, but not CFZ, suppressed cell migration and invasion similarly to *GALNT14* depletion.

### Attenuation of the TGFβ gene response by BTZ treatment or *GALNT14* knockdown

We hypothesized that a subset of the 101 genes down-regulated by both BTZ treatment and *GALNT14* depletion could account for anti-migration/invasion effects of BTZ. To investigate the drug’s mode of action, we conducted pathway enrichment analysis of the 101 genes and investigated the clinical significance of each pathway (Figure. 4A). Among the most enriched pathways was TGFβ signaling (hazard ratio [HR] = 1.2). BTZ treatment induced changes in expression of individual TGFβ signaling genes that were very similar to those induced by sh*GAL* (Figure. S4A for BTZ and Figure. S4B for *shGAL). Moreover, genes commonly down-regulated (e.g., INHBA, FST, and BMPR) among targets of TGFβ signaling were indeed suppressed (Figure. 4B)*. Suppression of the TGFβ-dependent gene signature, a common effect of BTZ treatment and *GALN14* depletion, was validated by reporter assays using the Smad-binding element (SBE), activin-response element (ARE), and BMP-response element (BRE) (Figure. 4C). Similarly, TGFβ reporter activity decreased after treatment with BTZ, but not CFZ (Figure. 4D), while β-catenin and Cyclin D1 were stabilized by proteasome inhibition following treatment with either BTZ or CFZ (Figure. S4C). Receptors activation by TGFβ transduce signal through direct phosphorylation of R-SMAD, including SMAD2 or SMAD3, which leads to interaction to co-SMAD (i.e. SMAD4) for nuclear translocation (34). Concomitant with the reduction in SMAD2 phosphorylation with (Figure. 4E) or without TGFβ stimulation (Figure. S4C), SMAD4 nuclear translocation was inhibited significantly by BTZ treatment (Figure. 4F). Consistently, TGFβ dependent gene responses (determined by SBE or BRE) upon TGFβ stimulation, were significantly attenuated by BTZ treatment (Figure. S4D). As depletion of SMAD4 was sufficient to inhibit both migration and invasion (Figures. 4G and H), inhibition of SMAD2 phosphorylation (Figures. 4E and S4C) and subsequent delay of SMAD4 nuclear translocation (Figure. 4F) by BTZ would be a possible mode of action of BTZ. It is important to note that the TGFβ signaling pathway has been studied extensively as a tumor suppressor, a tumor promoter (35), and a promoter of metastasis (36). To determine whether the TGFβ signaling response is associated with *GALNT14*, we selected a set of TGFβ downstream targets (37) and examined their correlations with *GALNT14* expression and patient prognosis (Figure. 4I). Notably, SMAD4-dependent targets *PCDH7* and *LAMC2*, previously shown to induce metastasis (38, 39) or tumorigenicity (40, 41), were highly correlated with *GALNT14* (Figure. 4I). Moreover, the SMAD4-dependent TGFβ targets was associated to RFS in the *GALNT14*-high group (P = 0.045, Figure. 4J), suggesting that some SMAD4-dependent targets responsible for cancer recurrence are strongly associated with *GALNT14* expression. These results imply that BTZ treatment, like *GALNT14* depletion, exerts its anti-metastatic effect by interfering with nuclear translocation of SMAD4 (Figure. 4F) and with the SMAD4-dependent gene expression response (Figure. 4D). Finally, we defined a set of genes commonly down-regulated by both BTZ and shGAL among the SMAD4 dependent targets as ‘*GALNT14*-TGFβ signature’ (See Materials and Methods). The average activity of the *GALNT14*-TGFβ signature strongly discriminated patient RFS (*P* = 4.0 × 10^−4^, Figure. 4K), and *GALNT14* expression was significantly higher in lung cancer patients with higher levels of the signature (*P* =0.025, Figure. 4L). Overall, these results suggest that suppressing TGFβ signaling and gene expression responses relevant to BTZ (similar to the response observed after *GALNT14* depletion) makes a major contribution to reducing migration and invasion.

**Figure 4.**
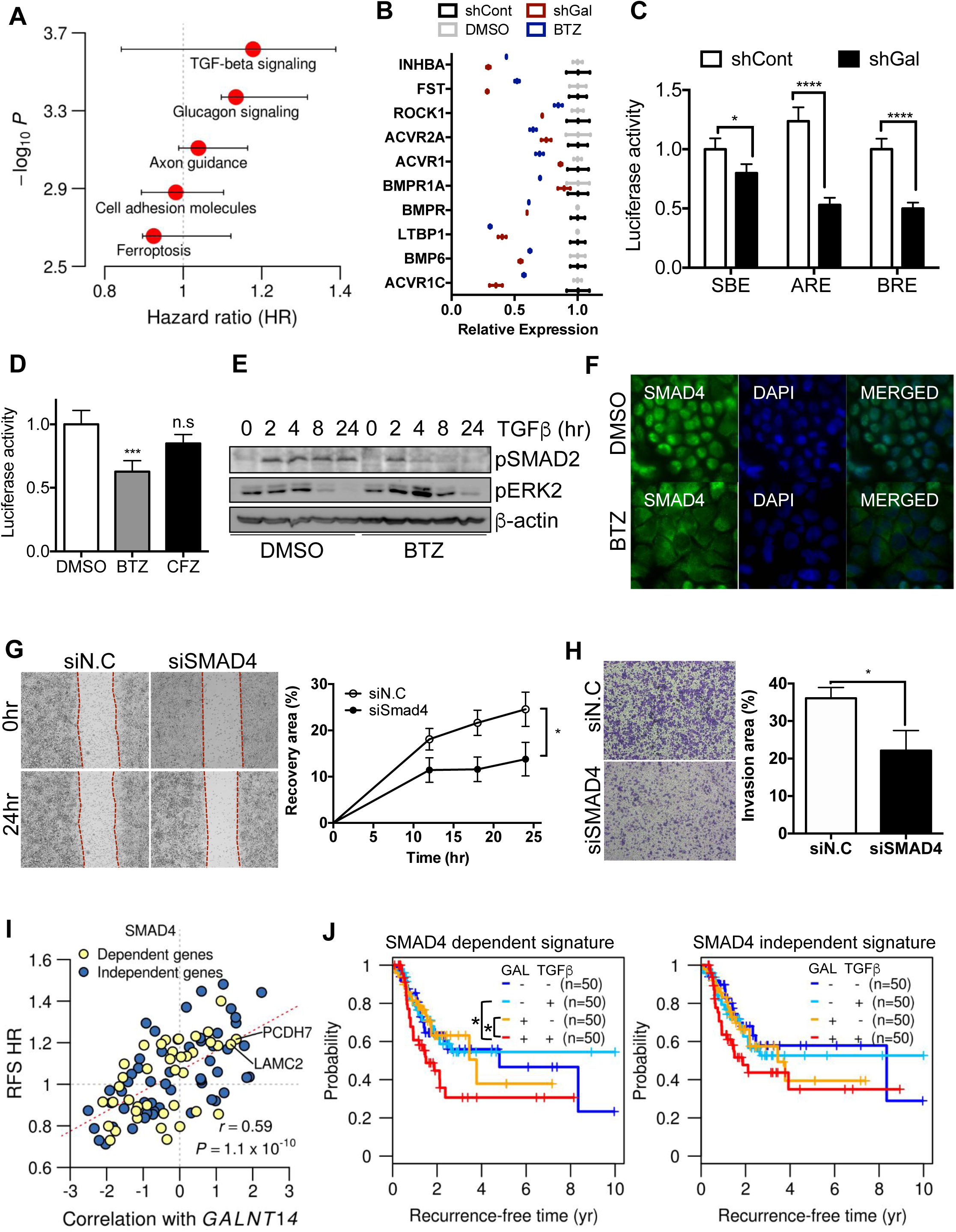
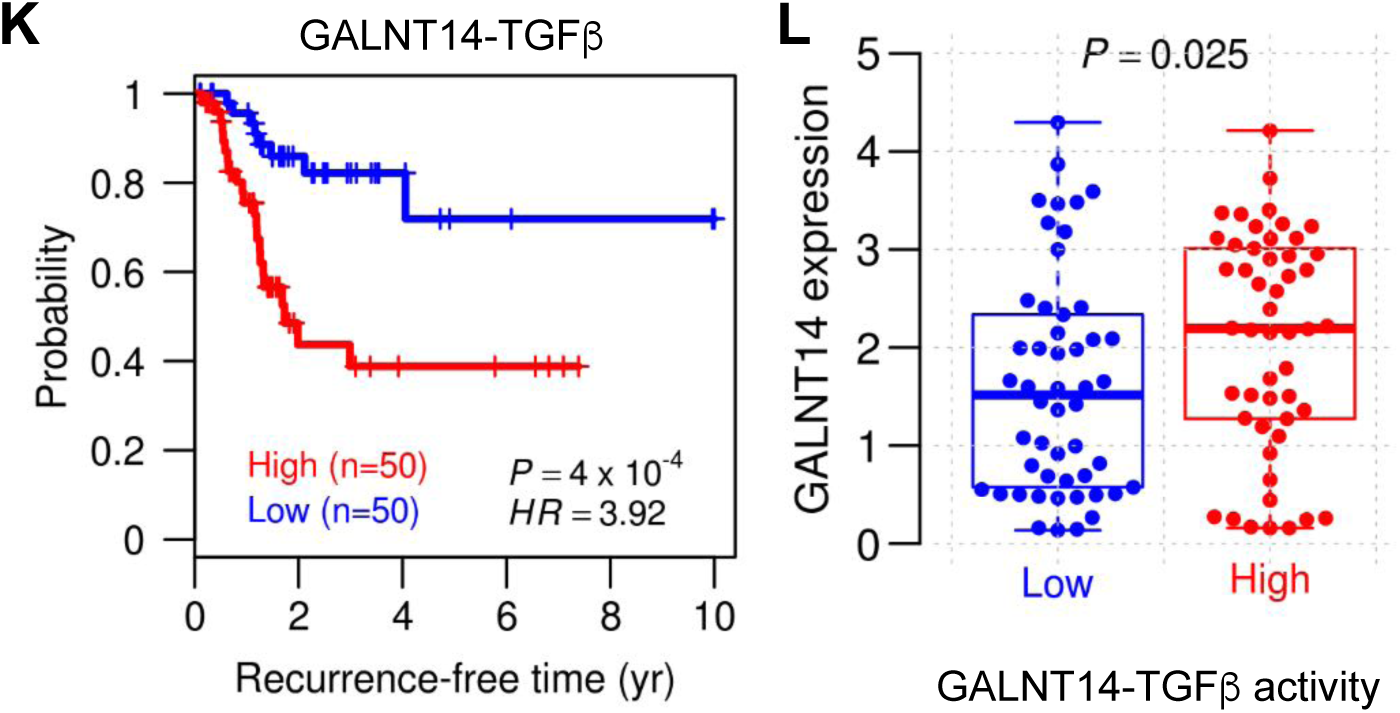
Attenuation of the TGFβ gene response by BTZ treatment or *GALNT14* knockdown. **A.** Enriched pathways in the 101 genes common to the down-regulated signatures by BTZ treatment and *GALNT14* knockdown (hypergeometric test, P value < 0.01 and FDR < 0.1), and the distribution of RFS HR for the pathway genes in TCGA LUAD cohort (red circle: the median HR of the pathway genes, horizontal bar: the range of HR in each pathway). **B.** mRNA expression of TGF-beta signaling genes among the 101 common signature genes by *GALNT14* depletion and BTZ treatment. **C.** Relative luciferase reporter activity (SBE, ARE and BRE) of shRNA control (shCont: white box) and *GALNT14* knockdown (shGal: black box) H460 cells was graphically presented. **D.** BRE luciferase activity at 24 hours after 20 nM of BTZ or CFZ treatment (n.s: not significant) **E.** Immunoblotting analysis for phosphorylated SMAD2 (pSMAD2) and phosphorylated ERK2 (pERK2) pretreated with DMSO or 20 nM of BTZ at indicated time after TGFβ (10 ng/ml) treatment, β-actin used as an equal protein loading control **F.** Fluorescent microscopic images for SMAD4 (green) after 20nM of BTZ treatment, DAPI (blue) for nuclear counterstaining **G-H** Representative microscopic images of cell migration (G) and invaded cells of two-chamber invasion assay (H) at 48 hours after introduction of control siRNA (siN.C) or siRNA for SMAD4 (siSMAD4) (left), Relative recovery ratio (G) or invaded area (H) was graphically presented (right). **I-L.** Analysis of TCGA LUAD cohort, **I.** The relationships between correlation score with *GALNT14* expression (z-transform of Spearman correlation, x-axis) and RFS HR (y-axis) of each gene in TGFβ target genes. The dependency of TGFβ target genes on Smad4 was labeled with two different colors (yellow: Smad4-dependent, blue: Smad4-independent). **J.** KM plot of RFS stratified by the combination of *GALNT14* expression (low and high) and TGFβ target activity (low and high). TGFβ target activity was measured using SMAD4 dependent genes (left) and independent genes (right) individually. Significant differences between the patient groups were marked with the asterisk (*). **K.** KM plot of RFS stratified by the *GALNT14*-TGFβ signature. *P* values and HR were calculated with the log-rank test and Cox regression, respectively. **L.** Differences in *GALNT14* expression by *GALNT14*-TGFβ signature in TCGA LUAD cohort. *P* values were calculated with Student’s t-test.

### *In vivo* validation of ant-metastatic effect of BTZ

Given the anti-migration/invasion effect of BTZ in a lung cancer cell model (Figure. 2), we next tested the *in vivo* efficacy of BTZ against cancer metastasis *in vivo*. For this purpose, local metastasis was induced in mice by tail vein injection of H460 lung cancer cells, followed by treatment with or without BTZ or CFZ twice weekly for three weeks (Figure. 5A). The proteasome-inhibitory effect of BTZ (0.1mg/kg) with CFZ (0.5 mg/kg) was examined by measuring proteasome activity in blood (Figure. 5B). Under this concentration, the mice tolerated both BTZ and CFZ, exhibiting neither significant loss of body weight nor any other abnormalities (Figure. 5C). Consistent with the *in vitro* assay, the number of metastatic nodules in the lungs of BTZ-treated mice was significantly lower than that in CFZ- or vehicle-treated animals (Figure. 5D, S5A and S5B). Close examination of cancer tissue also revealed that inflammatory lesions, which provide favorable microenvironments for tumor formation (42), were present in both CFZ- and vehicle-treated mice (Figure. S5C). These results were validated with further experiment to support the efficacy of BTZ on metastasis inhibition. BTZ treatment considerably attenuated lung colonization (Figure. 5E and S5D) in the presence of a proteasome inhibitory effect of BTZ (Figure. 5F) but no physiological abnormality (Figure. S5E). Taken together, the *in vivo* and *in vitro* data reveal that BTZ has a significant therapeutic advantage over CFZ in that it inhibits cancer metastasis without significant undesirable side effects.

**Figure 5.**
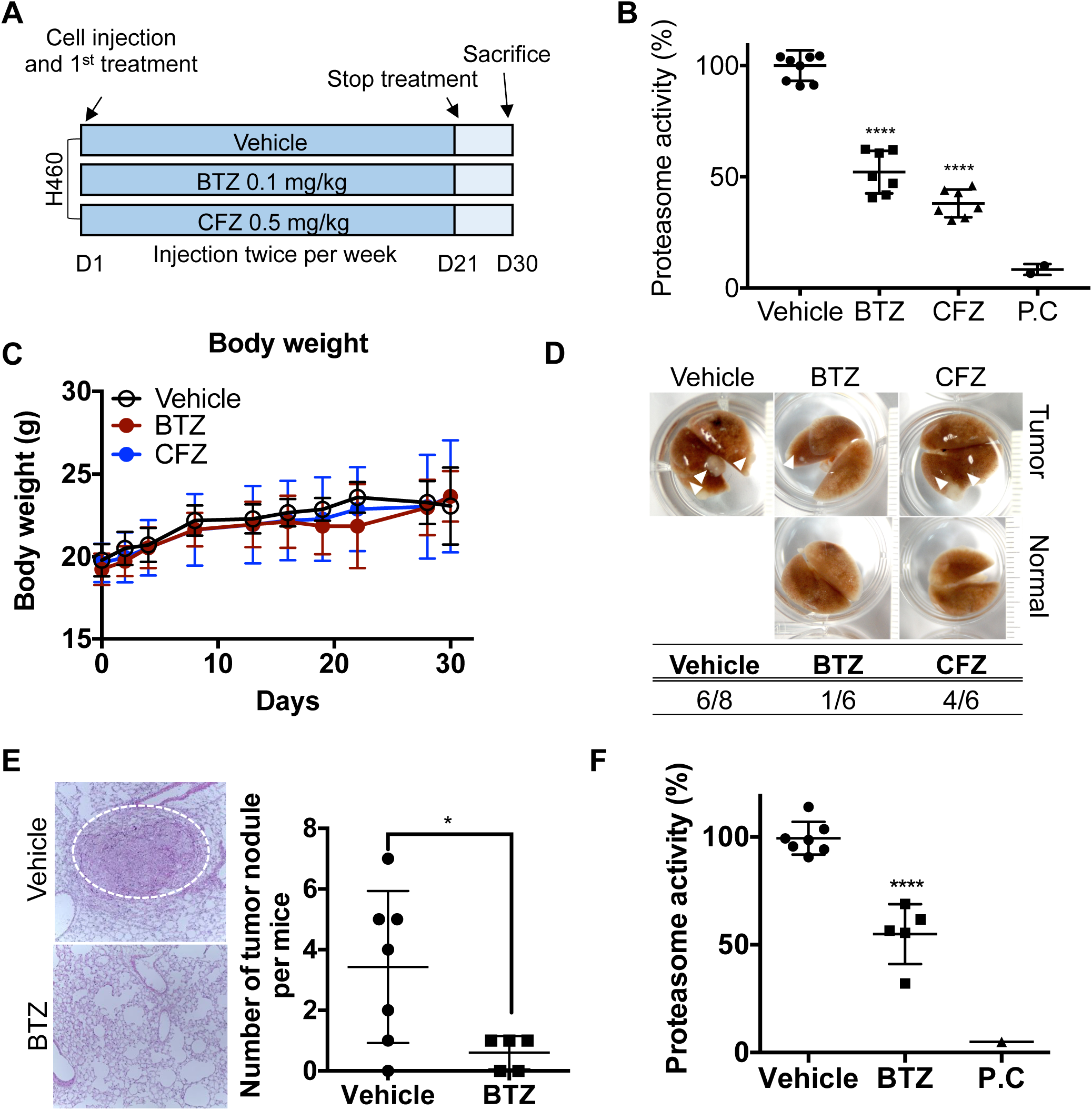
*In vivo* validation of anti-metastatic effect of BTZ. **A.** Schematic overview of *in vivo* experimental procedure **B-D.** Lateral tail vein injection of vehicle (saline, n=9) or BTZ (0.1 mg/kg, n=7) or CFZ (0.5 mg/kg, n=7) followed by H460 cells (B) Proteasome activity in whole blood collected 1 hour after BTZ or CFZ was graphically presented. (P.C: positive control, described in material method) (C) Body weight of each mouse monitored at indicated days, was graphically presented (D) Representative images of whole lung of each group was presented (top). White arrowheads indicate tumor nodule. Number of mice bearing tumor nodule was shown in the table (bottom). **E-F** Lateral tail vein injection of vehicle (saline, n=7) or BTZ (0.1 mg/kg, n=5) followed by H460 cells (E) Representative microscopic images of lung tumor bearing lesion (left) and graphical presentation of the number of metastatic tumor nodule (right) (F) Proteasome activity in whole blood collected 1 hour after BTZ was graphically presented. (P.C: positive control, described in material method)

Although many molecular targets for tumorigenesis and metastasis have been identified, most remain undruggable. For example, the GALNTs, expression of which is strongly associated with various properties of cancer (20–25), have yet to be drugged, although a few attempts have been made to develop inhibitors of GALNT-dependent O-linked glycosylation (29, 30). Thereby, instead of searching direct inhibitors of GALNT14, we adopted an *in silico* DR approach to reverse the *GALNT14-dependent* metastatic expression signature with the goal of finding a candidate drug that could interfere with the GALNT14-dependent cancer phenotype. Notably, *HOXB9* and *SOX4*, transcription factor genes regulated by *GALNT14*, are responsible for metastasis (23) and self-renewal (20), respectively, suggesting that downstream transcriptional modulation would be a promising strategy.

Unlike similar studies in the past that used the CMap method to analyze differences in gene expression signatures between normal and cancerous tissue (14, 43), we focused exclusively on genes (e.g., *GALNT14*) related to the pertinent phenotype (*i.e*., the metastatic gene signature) (Figure. 1) and identified BTZ as a drug candidate with novel anti-metastatic effects both *in vitro* (Figure. 2C–2H) and *in vivo* (Figure. 5). Importantly, we also demonstrated that the anti-metastatic effect of BTZ, in contrast to that of CFZ, was independent of proteasome inhibition (Figure. 3). Moreover, the metastatic gene signature of CFZ was also distinct from that of BTZ, whereas the proteasome gene signatures of the two drugs were relatively similar (Figure. 3). Recent studies examined the inhibitory effect of BTZ on TGFβ-dependent responses such as fibrosis (44) and survival of lymphoma (45), and the results support our conclusions. We also identified the *GALNT14*-TGFβ signature that serves as clear indicators of poor prognosis (Figure. 4). Thus, attenuation of TGFβ signaling by BTZ, depletion of *GALNT14*, or inhibition of TGFβ signaling all decrease invasive properties *in vitro* (Figure. 4) and lung metastasis *in vivo* (Figure. 5), suggesting the drug’s mode of action. Accordingly, the *GALNT14*-TGFβ signature represents potentially useful prognostic marker for lung cancer patients, and could be used alongside previously reported marker [*e.g*., the TGFβ-response signature (TBRS) (46) and the MAPK pathway activity score (MPAS) (47)]. Of note, although BTZ had no undesirable side effects in our *in vivo* experiments, the risk of peripheral neuropathy in patients treated with BTZ (48) merits a further search for other candidate drugs with safer profiles that could also reverse the gene signature associated with *GALNT14* or *GALNT14*-TGFβ activity. The continued search for improved candidates could be performed using recently reported *in silico* (or computational) DR tools (8, 49, 50). Notably in this regard, a recent *in silico* approach can predict candidate drugs capable of modulating the activities of oncogenic transcription factors, a class of proteins that has yet to be drugged (7). In the future, it would be interesting to apply this type of approach to modulate *GALNT14-regulated* transcription factor genes such as *HOXB9* and *SOX4*, which mediate metastasis (23) and self-renewal (20), respectively. Besides predicting potential candidate drugs, our *in silico* DR approach enabled identification of several marker genes that turned out to be strongly associated with clinical outcomes such as RFS. This strategy, which integrated multiple independent expression signatures from cancer patients, genetic perturbation (*e.g*., knockdown or overexpression), and drug treatment (CMap), would be applicable generally to any other types of target. Thus, our results provide a strong “proof-of-concept” that our DR method is a viable strategy for accessing undruggable molecular targets, leading to identification of candidate drugs that target specific cellular processes such as cancer metastasis.

## Methods

### Cell line establishment and Cell culture

H460, A549 and H1299 cell line which was purchased from Korean cell line bank (KCLB) were maintained in Dulbecco’s modified Eagle’s medium (DMEM), supplemented with 10% fetal bovine serum (FBS), gentamicin (50 μg/ml) at 37°C in a humidified atmosphere of 5% CO_2_ in the air. *GALNT14* knockdown cell lines with shRNA were established as previously described (23).

### Reagents and antibodies

The primary antibodies against cleavage caspase-3 (#9664), cleavage caspase-9 (#9505) and pSmad2 (#310S) were obtained from Cell Signaling Technology. Antibodies against β-Actin (sc-47778), PARP (sc-7150), p53 (sc-126), p27 (sc-528), p21 (sc-397), CyclinD1 (sc-718) and CyclinB1 (sc-245) were obtained from Santa Cruz Biotechnology Inc. and β-catenin (BD 610153) was purchased from BD Biosciences pharmigen. Bortezomib (S1013) and Carfilzomib (S2853) were purchased from selleckchem.

### RNA-sequencing and analysis

H460 cells treated with BTZ, CFZ, depleted of *GALNT14* (shGAL) and their control (DMSO, shCont) were prepared for RNA sequencing. Total RNA was isolated using the Trizol according to the manufacture instruction. For library construction, we used the TruSeq Stranded mRNA Library Prep Kit (Illumina, San Diego, CA). Briefly, the steps of strand-specific protocol are: first strand cDNA synthesis; second strand synthesized using dUTPs instead of dTTPs; end repair, A-tailing, and adaptor ligation; PCR amplification. Then, each library was diluted to 8 pM for 76 cycles of paired-read sequencing (2 × 75bp) on the Illumina NextSeq 500 per the manufacturer’s recommended protocol. Read sequences were aligned to the reference genome (UCSC hg19) and the mapped counts per gene were quantified by STAR (51). The raw counts were normalized to CPM (counts per million) based on the trimmed mean of M-values (TMM) normalization method using R package ‘edgeR’. Differential expression analyses between samples (shCont vs shGAL, DMSO vs BTZ, or DMSO vs CFZ) were performed through DESeq2 (52).

### TCGA data processing and analysis

TCGA lung adenocarcinoma (LUAD) cohort (n=576), containing mRNA gene expression and clinical data on 388 primary lung cancers, 128 lung cancers with recurrence, and 59 benign lung tissues were collected from the Broad GDAC Firehose (https://gdac.broadinstitute.org/). Total 494 patients with clinical information tracked for at least one month were used for survival analysis. For 20,531 genes, all patients were divided into high and low groups by the median expression of each gene and relapse-free survival analysis was performed according to the group difference. Hazard ratio and *P* value were calculated by Cox proportional hazards regression model and log-rank test respectively. With 58 patients who have gene expression profiles of normal benign tissues, differential expression in LUAD compared to matched normal samples were measured from the likelihood ratio test. RNA-seq profiles were normalized and processed using R package ‘limma’ and ‘DESeq2’, and survival analysis was conducted by R package ‘survival’.

### Metastasis and Tumorigenesis signatures

From MSigDB manually curated gene sets (C2), we collected the 35 metastasis- and 44 tumor-related gene sets that are annotated by ‘metastasis’ / ‘epithelial-mesenchymal transition’ and ‘cancer’ / ‘tumorigenesis’, respectively. We took only Up-regulated genes when both UP- or DOWN-regulated sets were available. Then, we selected 32 and 23 genes as the metastatic and the tumorigenesis signature, respectively, by taking the consensus genes common to at least three or more gene sets.

### CMap analysis to predict candidate drugs

The updated CMap dataset, or LINCS L1000, provides an extensive catalog of transcriptome profiles for 71 human cell lines treated by 20,413 small molecules. We obtained the expression dataset (level 5, replicate-collapsed z-score) from Gene Expression Omnibus (GSE92742), each of which represents normalized genome-wide differential expression profiles under a unique experimental condition (cell line, drug, treated dose/time, batch). We used only the 60,321 profiles of high-quality (distil_cc_q75 ≧ 0.2, pct_self_rank_q25 ≦ 0.05, and distil_nsample ≧ 3) among the total 205,034 profiles. Unlike the original CMap based on microarray, a single drug may have multiple expression profiles depending on cell line, dose, time, and sample batch. Therefore, instead of using the original rank-based Kolmogorov-Smirnov (KS) test, we made a modified analytic scheme to handle such redundancy. For prediction of drugs mimicking the down-regulation of *GALNT14*-associated metastatic genes (i.e. *GALNT14* signature), we calculated Jaccard index between the *GALNT14* signature and the 100 most down-regulated genes of CMap profiles as similarity score (S). Then, the similarity scores of a drug were merged into a single DR score (drug repositioning score), which was calculated as the negative logarithm of the hypergeometric P-value for over-representation of its similarity scores within the top 10% of the total similarity scores.

### GALNT14-TGFβ signature

TGFβ downstream target genes (39 SMAD4 dependent genes and 65 SMAD4 independent genes) were collected from the literature (37). Among the SMAD4 dependent targets, a set of genes commonly down-regulated by both BTZ and shGAL (*ATF3, ARNTL, COPA, NEDD9, LAMC2, RAB27B*, and *PCDH7*) was defined as ‘*GALNT14*-TGFβ signature’. To measure the average activity of the signature, KS statistic was used to estimate the degree of up- or down-regulation of the seven genes in a sample’s gene expression profile. In TCGA LUAD cohort, the patients with activity in the top 10% (n=50) or lower 10% (n=50) were classified as ‘high’ or ‘low’ respectively.

### Statistical analysis

The graphical data were presented as mean ± S.E.M. Statistical significance among the three groups and between groups was determined using one-way or two-way analysis of variance (ANOVA) following Turkey post-test and Student’s t-test respectively. Significance was assumed for p < 0.05 (*), p < 0.01 (**), p < 0.001 (***).

## SUPPLEMENTARY DATA

Supplementary information is available from the Wiley Online Library or from the author.

## AUTHORS’ CONTRIBUTIONS

HJ.C and W.K conceived the overall study design and led the experiments. OS.K and H.L mainly conducted the experiments, data analysis, and critical discussion of the results. HJ.K, JE.P, and W.L conducted the mouse xenograft experiments. S.K, JH.K and M.K generated and analyzed RNAseq data. All authors contributed to manuscript writing and revising, and endorsed the final manuscript.

## ACKNOWLEDGEMENT

We appreciate Jeong-Hwan Kim, Seon-Young Kim and Dong-Uk Kim at Korea Research Institute of Bioscience and Biotechnology (KRIBB) for helpful discussions.

