## Supplementary Figures and legends for "Computational drug repositioning of bortezomib to reverse metastatic effect of *GALNT14* in lung cancer"

Supplementary information for

### Computational drug repositioning identifies bortezomib as a novel metastatic inhibitor of lung cancer

Running title**: A data-driven route to drug discovery for undruggable targets**

This PDF file includes:

Supplementary Materials and Methods

Supplementary Figures 1-5

Supplementary Table 1

#### Supplementary Materials and Methods

##### Immunoblotting

Cells were lysed with Tissue lysis buffer or RIPA buffer with 10uM sodium vanadate and 1mM protease inhibitor (Roche) and immunoblotting was performed as described elsewhere.

##### Real-time PCR

Total RNA was extracted by using Trizol (Invitrogen), followed by RT-PCR to generate the first strand cDNA (Takara RR036A), and the cDNA was subjected to Real-time PCR (Roche LightCycler 480 II) with SYBR Green (Takara RR420).

##### Migration assay

For wound healing assay, after 24 hours incubation on serum starvation condition, the compact cell was scratched by using sterile micropipette tip with indicated drug treatment, Cell migration was monitored and live image was taken by using Juli stage for 48 hours. For coverslip migration assay, coverslip on which cells were plated compactly, was transferred to empty 6 well plate as described previously. The pixel count of an area of cell migration was obtained with Image J software.

##### Trans-well invasion assay

Trans-wells (6.5 mm) with 8uM pore polycarbonate membrane insert (Corning, #3422) was embedded with matrigel (BD Bioscience) and bottom of the Matrigel-embedded insert was coated with 0.2% gelatin. Drug pretreated cell was allowed to invade through the Matrigel-embedded barrier. After 24 hours' incubation, the membrane was fixed with 4% formaldehyde and stained with 0.1% crystal violet. The images were taken with a light microscope imaging system (ProgRes C3)

**FACS analysis**

FACS Caliber (BD Bioscience) was used for Annexin V staining and PI staining. For Annexin V staining, drug pretreated cells were washed twice with PBS and stained with FITC-Annexin V and 7-AAD for 30 minutes at RT in dark. For cell cycle analysis, cells were washed twice with PBS and stained with 1ug/ml propidium iodide in the presence of 500ug/ml RNase. All the FACS analysis were performed in accordance with manufacturer's instruction

**Dual luciferase assay**

Cells were transfected with SBE, ARE and BRE luciferase vector and pRL plasmid with Lipofectamine 2000. After transfection, reporter assays were performed according to with manufacturer's instructions (Promega).

**Animal model for tumorigenesis and metastasis**

All experiments were performed according to protocols approved by the Institutional Animal Care and Use committees of Seoul National University (SNU-180717-1). Female BALB/c nude mice were purchased at 4 weeks of age. For the experimental tumorigenesis assay, cells ($1x10^7$) were subcutaneously injected at both right and left flanks of nude mice. For the experimental lung metastasis assay, cells ($2x10^6$) were injected into lateral tail veins of mice. Bortezomib and carfilzomib were also injected via tail vein into mice bearing H460 at the dose of 0.1 mg/kg and 0.5 mg/kg respectively. Blood sample were collected from the retro-orbital plexus of the mice with microhematocrit tube at 1hour post-dosing. Mice were sacrificed at 4-6weeks after cancer cell injection, and section of lung was stained with H&E.

**Proteasome activity assay**

Cell lysate or blood sample was 9X diluted with passive lysis buffer (Promega, Madison, WI, USA). The diluted samples in assay buffer (20 mM Tris-Cl pH 8.0 and 500 μM EDTA) were incubated with 100 mM Suc-LLVY-AMC (Bachem, Bubendorf, Switzerland). The proteasome activity was determined by monitoring the cleavage rate of AMC. Fluorescence signals were detected using a SpectraMax M3 microplate reader (Molecular Devices, CA, USA). Positive control (P.C) was used to ensure the complete proteasomal inhibition, which was prepared by the blood samples from vehicle mice incubated with 1 μM of bortezomib 1hour prior to the addition of Suc-LLVY-AMC.

**Supplementary Figure legends**

**Figure S1 A**. Kaplan–Meier (KM) plot of overall survival in LUAD patients stratified by individual gene expression of seven candidate biomarkers. **B**. Distributions of dependency scores of each gene according to cell lines from metastatic lung cancer, and primary lung cancer. The dependency scores indicating genetic vulnerability to each of a broad range of cancer cell lines were obtained from the *Project Achilles* data. **C.** *GALNT14* expression in normal and tumor samples of 15 TCGA cohorts in which more than 10 normal (blood or adjacent tissue) samples are available (Table S1).

**Figure S2 A**. List of top 10 candidate drugs predicted by using two *GALNT14* signatures (TCGA: left table and H460: right). **B.** Venn diagram of the number of differentially down-regulated genes by BTZ, DEX, and depletion of *GALNT14* (sh*Gal*). Representative genes significantly down-regulated were listed on the Venn diagram. **C.** mRNA levels of *GALNT14* and representative genes (*SOX4, AREG, VCAN, HOXB9* and *FST*) in shCont and shGal **D.** mRNA expression of *SOX4* and *VCAN* after treatment of BTZ and DEX with indicated concentration was graphically presented. **E.** Immunoblot for β-catenin, Cyclin B1, p21 and p53 after treatment of BTZ with indicated concentration. β-actin was used for loading control. **F.** Cell migration rate from the coverslip to the empty area was examined after treatment of BTZ at the indicated concentration. Total pixel counts of the area of cell migration were quantified as a graph. **G.** mRNA expression of *GALNT14* in *GALNT14* knockdown cell line (sh*Gal*) and BTZ treatment compared to control (shCont or DMSO). **H.** Immunoblotting for apoptosis marker such as cleaved PARP (F: Full length, T: Truncated), cleaved caspase 3 (cCASP3) and cleaved caspase 9 (cCASP9) after BTZ treatment at indicative concentration, β-actin and Cyclin D1 was used for control of an equal protein loading and inhibition of proteasome activity respectively. **I.** Cell proliferate rate was measured with the indicated dose of BTZ **J.** mRNA expression (up) and immunoblotting analysis (bottom) for GALNT14 from indicated lung cancer cell lines (A549, H1299, and H460), β-actin was used for loading control. **K-L.** Relative recovery ratio from wound healing assay (**K**) and invaded area from two chamber invasion assay (**L**) after BTZ treatment (20nM) in A549 (left) and H1299 (right) was graphically presented (n.s: not significant). **M.** Immunoblot for p53 and p27 or Cyclin D1 after a various dose of BTZ treatment in A549 (top) and H1299 (bottom), β-actin was used as an equal loading control.

**Figure S3 A**. Proteasome activity was measured after the indicated dose of CFZ treatment **B.** Representative microscopic images of cell migration at 43 hours after 20nM of CFZ and IXZ treatment (left), relative recovery ratio was graphically presented (right, n.s: not significant)

**Figure S4 A-B.** TGFβ signaling pathway that reflects changes (z-score) in gene expression due to BTZ treatment (A) and *GALNT14* depletion (B) in H460. Annotations on nodes and edges were obtained from KEGG and the network was rendered by software Cytoscape. **C.** Immunoblotting for β-catenin, CyclinD1 and pSmad2 after BTZ (20nM) and CFZ (20nM) treatment, β-actin used as an equal protein loading control **D.** Luciferase activity of SBE and BRE with indicated time with TGFβ (10ng/ml) stimulation in either DMSO or BTZ (20nM) pretreated H460 cell.

**Figure S5 A.** Gross picture of lungs in each group of condition **B-C.** Representative microscopic images of H&E staining of mouse lung tissues with (B) tumor-bearing mice after treatment of BTZ (0.1mg/kg) and CFZ (0.5mg/kg), with (C) inflammatory lesions (indicated by a white arrow) from vehicle and CFZ treated mice **D.** Microscopic images of H&E staining of lung section from all tumor-bearing mice after treatment of vehicle and BTZ (0.1mg/kg), black arrowheads indicate lesion of metastatic tumor nodule. **E.** Graphical presentation of total body weight of each mouse at indicated days after vehicle or BTZ (0.1mg/kg) treatment

**Figure. S1**

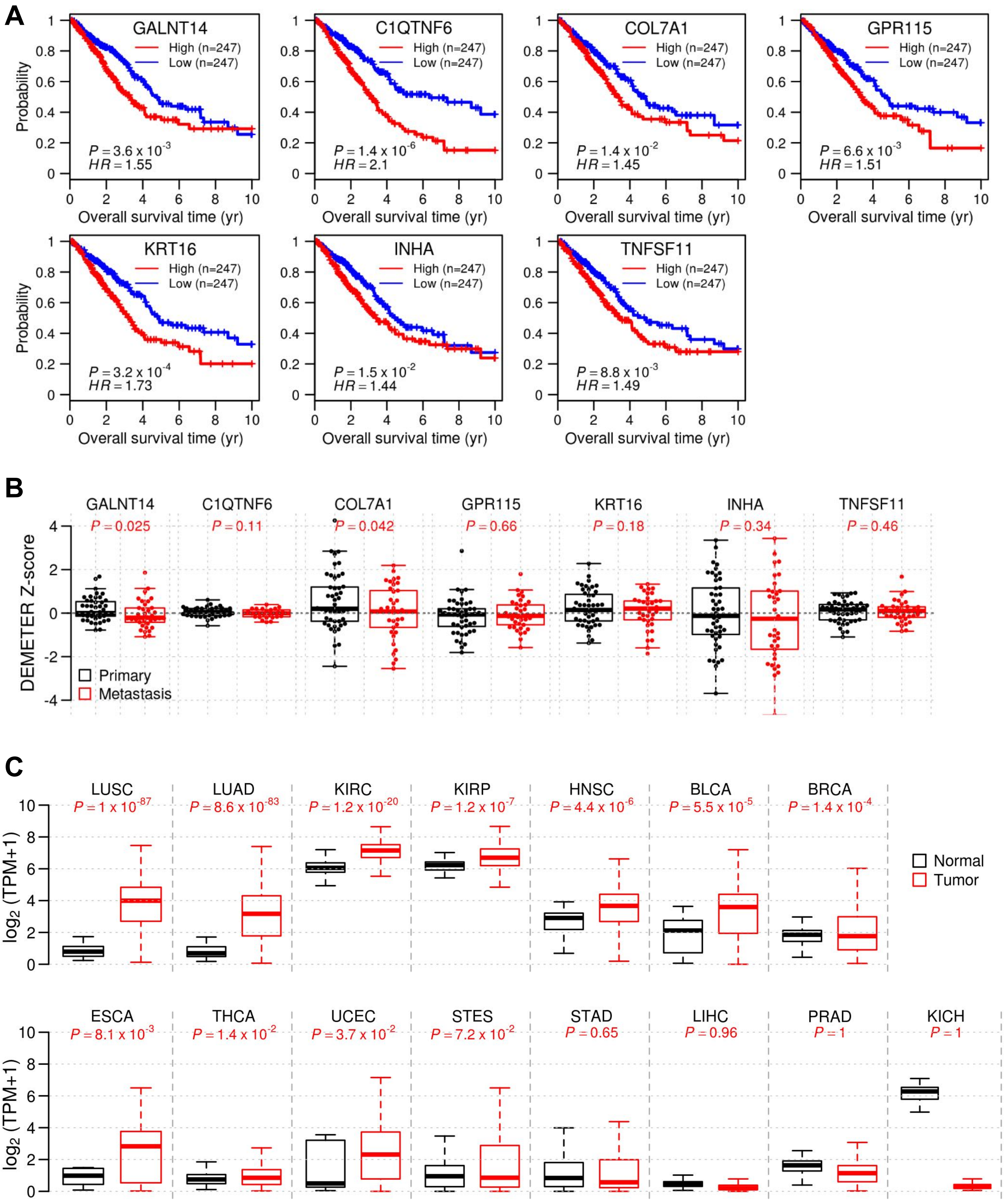
A
GALNT14
C1QTNF6
COL7A1
GPR115
KRT16
INHA
TNFSF11
High (n=247)
Low (n=247)
Probability
Overall survival time (yr)
P = 3.6 x 10^-3 HR = 1.55
P = 1.4 x 10^-6 HR = 2.1
P = 1.4 x 10^-2 HR = 1.45
P = 6.6 x 10^-3 HR = 1.51
P = 3.2 x 10^-4 HR = 1.73
P = 1.5 x 10^-2 HR = 1.44
P = 8.8 x 10^-3 HR = 1.49
B
P = 0.025
P = 0.11
P = 0.042
P = 0.66
P = 0.18
P = 0.34
P = 0.46
DEMETER Z-score
Primary
Metastasis
C
LUSC P = 1 x 10^-87
LUAD P = 8.6 x 10^-83
KIRC P = 1.2 x 10^-20
KIRP P = 1.2 x 10^-7
HNSC P = 4.4 x 10^-6
BLCA P = 5.5 x 10^-5
BRCA P = 1.4 x 10^-4
ESCA P = 8.1 x 10^-3
THCA P = 1.4 x 10^-2
UCEC P = 3.7 x 10^-2
STES P = 7.2 x 10^-2
STAD P = 0.65
LIHC P = 0.96
PRAD P = 1
KICH P = 1
log2 (TPM+1)
Normal
Tumor

### Figure. S2

**A**

| Rank | Name | DRscore |
|---|---|---|
| 1 | Vemurafenib | 3 |
| 2 | Thalidomide | 2.36 |
| 3 | Estradiol | 2.26 |
| 4 | Diltiazem | 2.19 |
| 5 | Chloroquine | 2.13 |
| 6 | Lapatinib | 2.04 |
| 7 | Dexamethasone | 1.98 |
| 8 | Bortezomib | 1.92 |
| 9 | Triamcinolone | 1.84 |
| 10 | Pazopanib | 1.81 |

| Rank | Name | DRscore |
|---|---|---|
| 1 | Domperidone | 2.9 |
| 2 | Dexamethasone | |
| 3 | Mosapride | 2.48 |
| 4 | Terbutaline | 2.43 |
| 5 | Gefitinib | 2.4 |
| 6 | Topiramate | 2 |
| 7 | Bortezomib | 1.92 |
| 8 | Dextromethorphan | 1.89 |
| 9 | Timolol | 1.89 |
| 10 | Hydrocortisone | 1.8 |

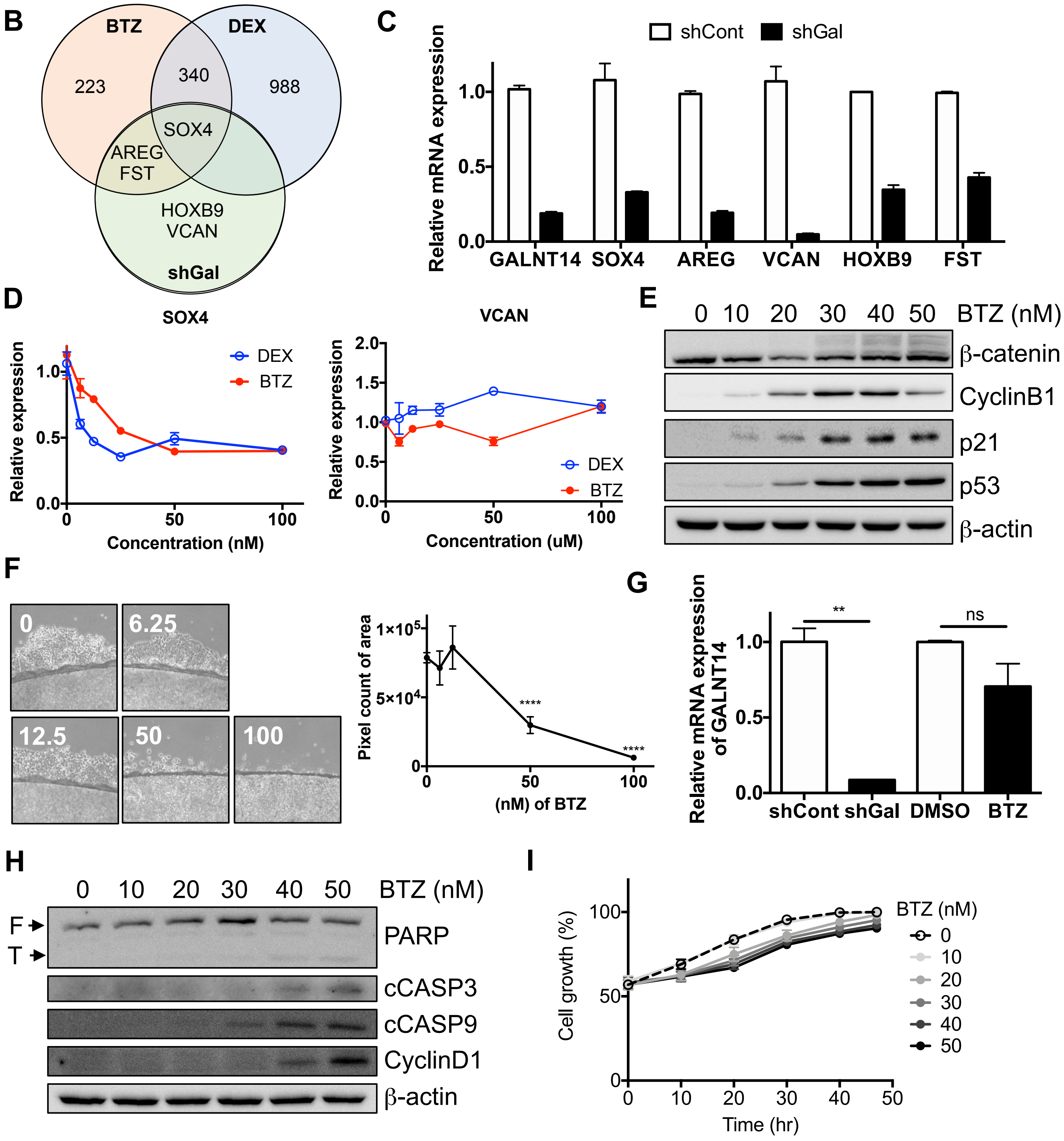

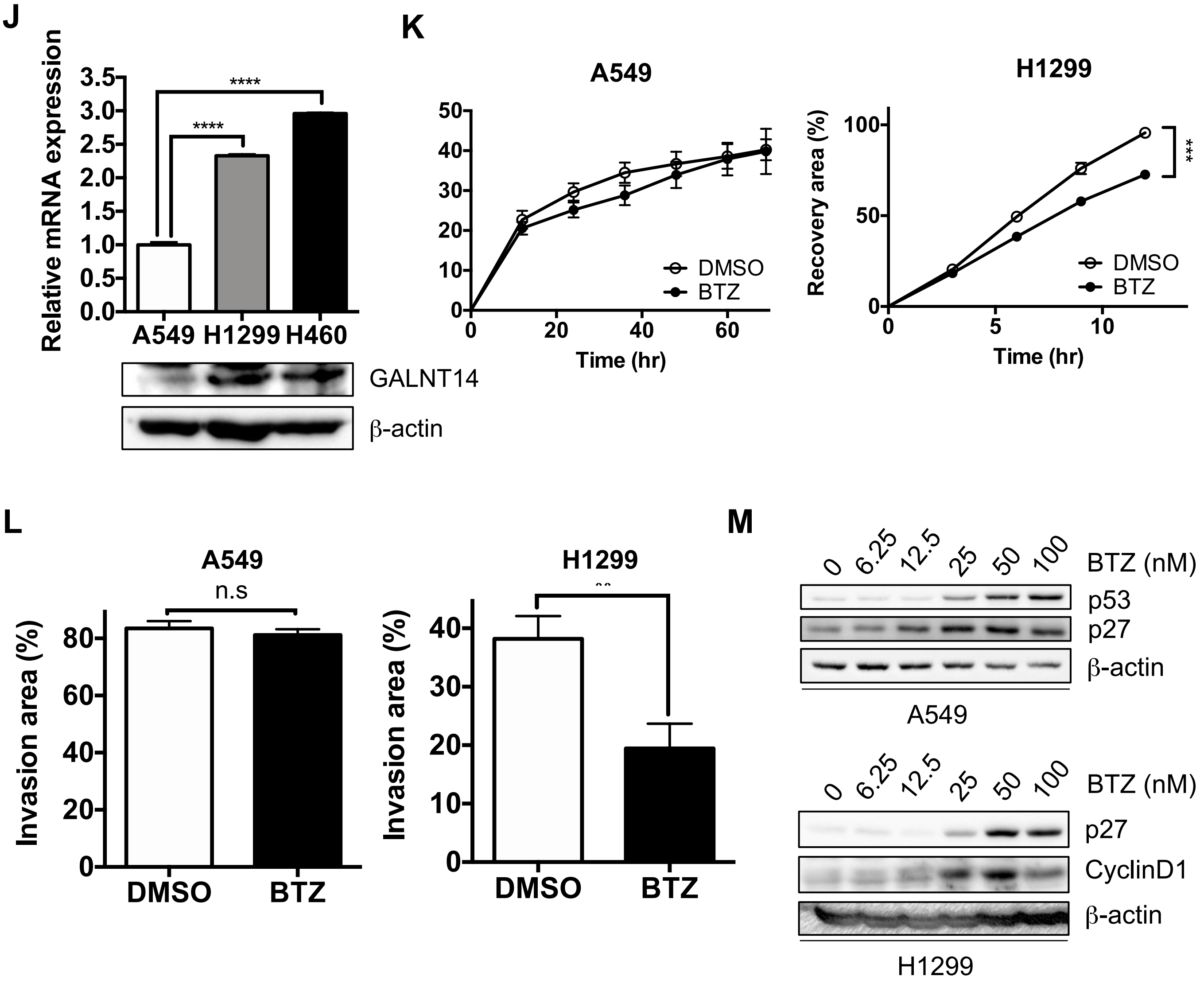
J
Relative mRNA expression
3.5
3.0
2.5
2.0
1.5
1.0
0.5
0.0
****
****
A549
H1299
H460
GALNT14
β-actin
K
A549
50
40
30
20
10
0
0
20
40
60
Time (hr)
DMSO
BTZ
H1299
Recovery area (%)
100
50
0
0
5
10
Time (hr)
***
DMSO
BTZ
L
A549
n.s
Invasion area (%)
80
60
40
20
0
DMSO
BTZ
H1299
Invasion area (%)
40
30
20
10
0
DMSO
BTZ
M
0
6.25
12.5
25
50
100
BTZ (nM)
p53
p27
β-actin
A549
0
6.25
12.5
25
50
100
BTZ (nM)
p27
CyclinD1
β-actin
H1299

Figure. S3

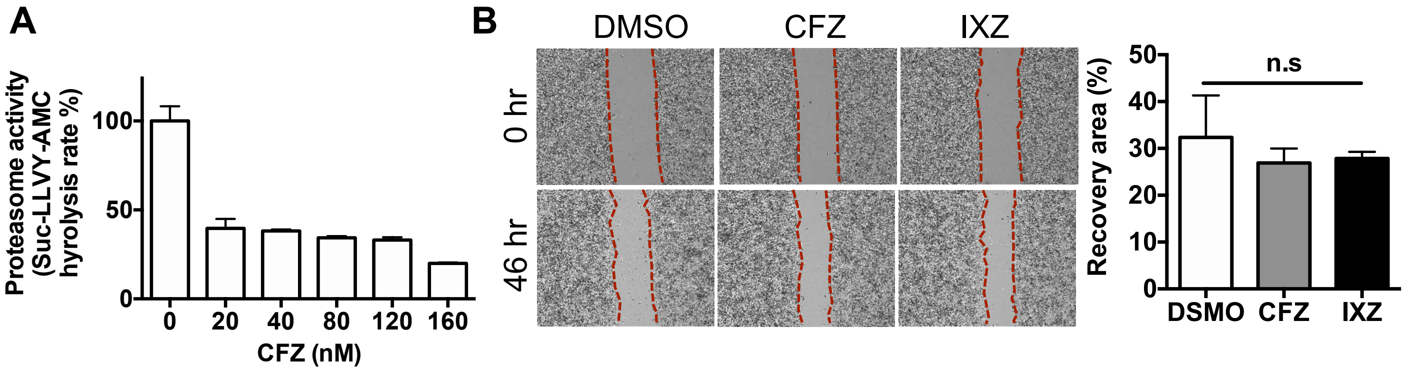
A
Proteasome activity
(Suc-LLVY-AMC
hyrolysis rate %)
100
50
0
0
20
40
80
120
160
CFZ (nM)
B
DMSO
CFZ
IXZ
0 hr
46 hr
50
40
30
20
10
0
Recovery area (%)
n.s
DSMO
CFZ
IXZ

Figure. S4

A *shGAL*

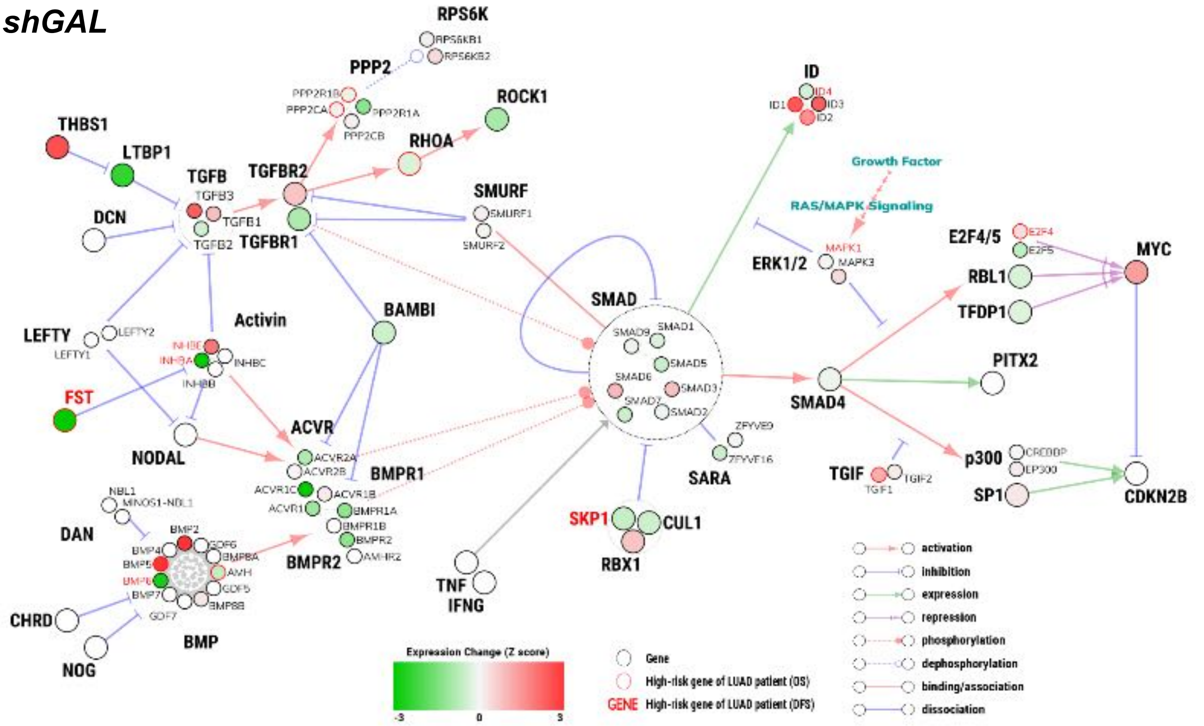

B BTZ

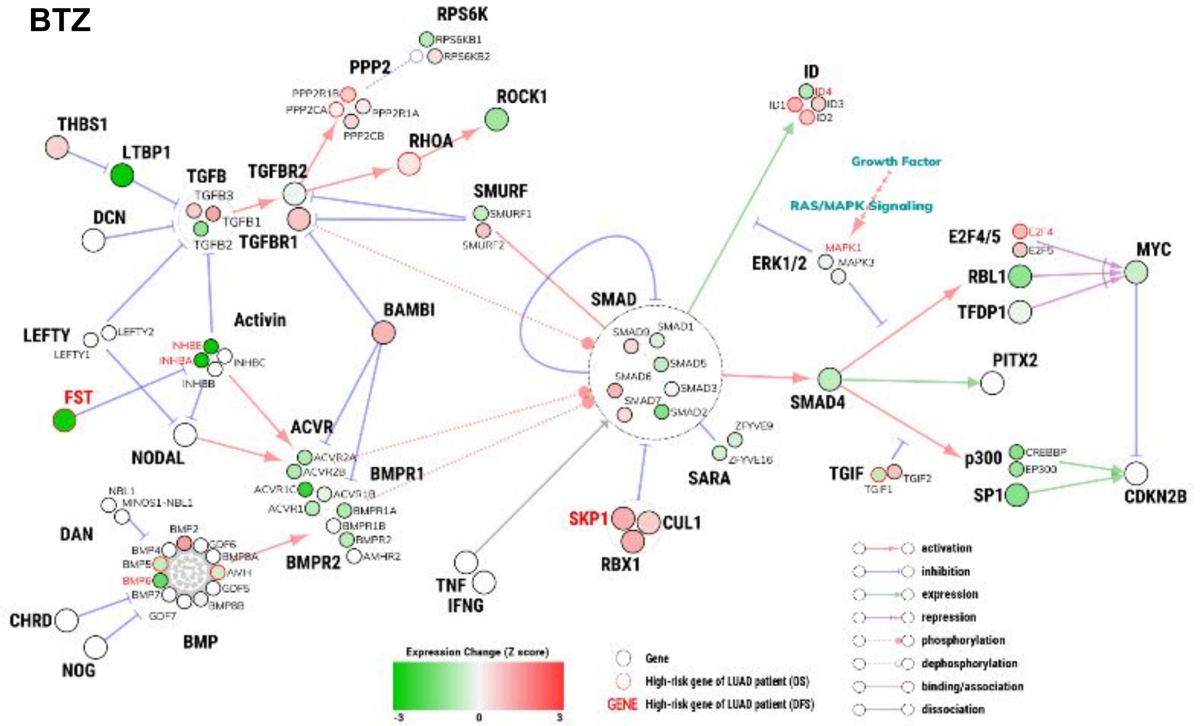

C

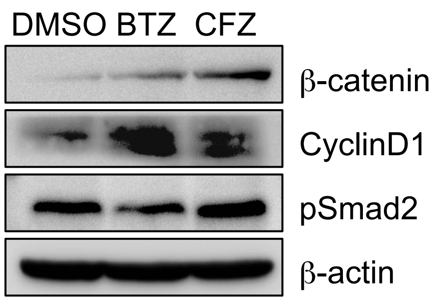


D

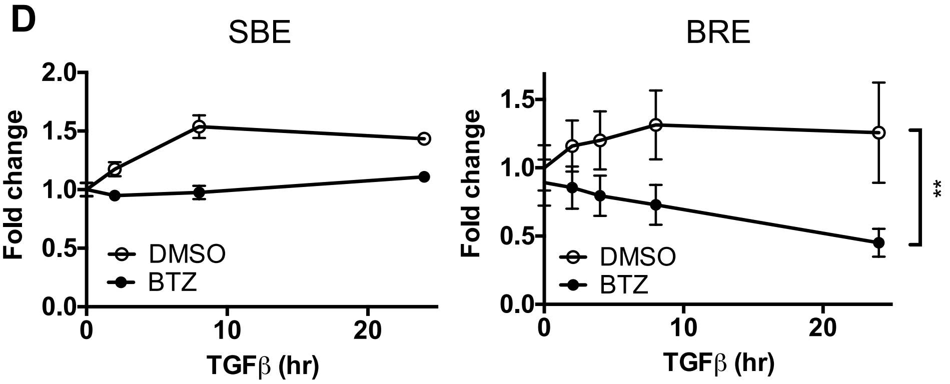

Figure. S5

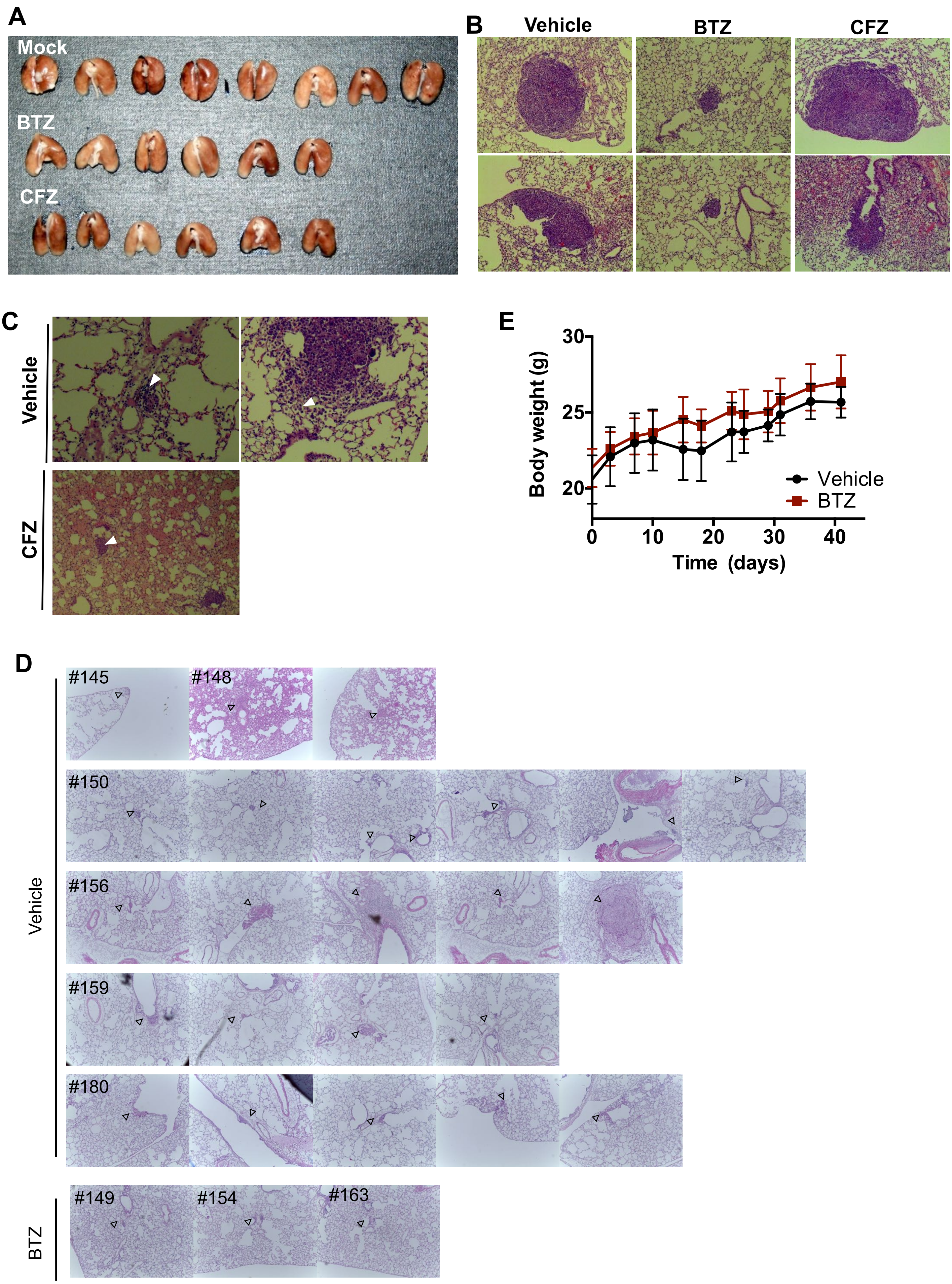
A
Mock
BTZ
CFZ
B
Vehicle
BTZ
CFZ
C
Vehicle
CFZ
E
Body weight (g)
30
25
20
0
10
20
30
40
Time (days)
Vehicle
BTZ
D
#145
#148
#150
#156
#159
#180
Vehicle
#149
#154
#163
BTZ

| Abbreviation | Cancer | # Normal sample | # Tumor sample | # Total sample |
|---|---|---|---|---|
| ACC | Adrenocortical carcinoma | 0 | 79 | 79 |
| BLCA | Bladder Urothelial Carcinoma | 19 | 408 | 427 |
| BRCA | Breast invasive carcinoma | 112 | 1100 | 1212 |
| CESC | Cervical squamous cell carcinoma and endocervical adenocarcinoma | 3 | 306 | 309 |
| CHOL | Cholangiocarcinoma | 9 | 36 | 45 |
| COAD | Colon adenocarcinoma | 0 | 191 | 191 |
| DLBC | Lymphoid Neoplasm Diffuse Large B-cell Lymphoma | 0 | 48 | 48 |
| ESCA | Esophageal carcinoma | 11 | 185 | 196 |
| GBM | Glioblastoma multiforme | 5 | 166 | 171 |
| HNSC | Head and Neck squamous cell carcinoma | 44 | 522 | 566 |
| KICH | Kidney Chromophobe | 25 | 66 | 91 |
| KIRC | Kidney renal clear cell carcinoma | 72 | 534 | 606 |
| KIRP | Kidney renal papillary cell carcinoma | 32 | 291 | 323 |
| LAML | Acute Myeloid Leukemia | 0 | 173 | 173 |
| LGG | Brain Lower Grade Glioma | 0 | 530 | 530 |
| LIHC | Liver hepatocellular carcinoma | 50 | 373 | 423 |
| LUAD | Lung adenocarcinoma | 59 | 517 | 576 |
| LUSC | Lung squamous cell carcinoma | 51 | 501 | 552 |
| MESO | Mesothelioma | 0 | 87 | 87 |
| OV | Ovarian serous cystadenocarcinoma | 0 | 307 | 307 |
| PAAD | Pancreatic adenocarcinoma | 4 | 179 | 183 |
| PCPG | Pheochromocytoma and Paraganglioma | 3 | 184 | 187 |
| PRAD | Prostate adenocarcinoma | 52 | 498 | 550 |
| READ | Rectum adenocarcinoma | 0 | 72 | 72 |
| SARC | Sarcoma | 2 | 263 | 265 |
| SKCM | Skin Cutaneous Melanoma | 1 | 472 | 473 |
| STAD | Stomach adenocarcinoma | 35 | 415 | 450 |
| STES | Esophagus-Stomach Cancers | 46 | 600 | 646 |
| TGCT | Testicular Germ Cell Tumors | 0 | 156 | 156 |
| THCA | Thyroid carcinoma | 59 | 509 | 568 |
| THYM | Thymoma | 2 | 120 | 122 |
| UCEC | Uterine Corpus Endometrial Carcinoma | 11 | 370 | 381 |
| UCS | Uterine Carcinosarcoma | 0 | 57 | 57 |
| UVM | Uveal Melanoma | 0 | 80 | 80 |
